# The CoREST complex is a therapeutic vulnerability in malignant peripheral nerve sheath tumors

**DOI:** 10.1101/2024.08.17.607802

**Authors:** Imad Soukar, Robert Fisher, Sanjana Bhagavatula, Marianne Collard, Philip A. Cole, Rhoda M. Alani

**Affiliations:** Department of Dermatology, Boston University Chobanian and Avedisian School of Medicine, Boston, Massachusetts, USA; Division of Genetics, Departments of Medicine and Biological Chemistry and Molecular Pharmacology, Harvard Medical School and Brigham and Women’s Hospital, Boston, Massachusetts, USA

**Keywords:** CoREST, Corin, HDAC, LSD1, MPNST, Epigenetics, NF1, PRC2, RNA-seq, ATAC-seq

## Abstract

Malignant peripheral nerve sheath tumor (MPNST) is a highly aggressive sarcoma that may be seen in patients with neurofibromatosis type 1 (NF1) or occur sporadically. While surgery is the primary treatment for localized MPNST with a 61.9% overall survival rate, metastatic disease is often fatal due to resistance to systemic therapies which underscores the urgent need for effective treatments. MPNSTs frequently harbor inactivating driver mutations in the PRC2 epigenetic repressor complex suggesting epigenetic therapies may represent a specific vulnerability in these tumors. Here, we investigate the role of the LSD1-HDAC1-CoREST (LHC) repressor complex in mediating MPNST tumor growth and progression. Our findings demonstrate that the LHC small molecule inhibitor, corin, induces apoptosis and significantly inhibits proliferation in MPNST cells. Transcriptomic analysis of corin-treated MPNST cells demonstrates specific increases in genes associated with axonogenesis and neuronal differentiation as well as altered extracellular matrix; additionally, corin treatment is shown to inhibit MPNST invasion in vitro. These results underscore the critical role of the LHC complex in facilitating MPNST growth and progression and suggest that targeting the LHC complex represents a promising therapeutic approach for this aggressive malignancy.

## Introduction

Malignant peripheral nerve sheath tumors (MPNSTs) are rare soft tissue sarcomas which are typically treated by surgical resection with a relatively high recurrence rate (Yao et al., 2023), and are frequently prone to metastasis (Acem et al., 2021). Patients with a germline mutation in the tumor suppressor gene, neurofibromatosis type 1 (NF1), carry a lifetime susceptibility to MPNST and account for over 50% of cases, with roughly 10% of NF1 patients developing MPNST (Knight et al., 2022), while sporadic MPNSTs may also carry NF1 mutations and occur at a rate of 0.001% (Cashen et al., 2004; Grobmyer et al., 2008; Gupta and Maniker, 2007). MPNSTs frequently possess inactivating mutations in genes associated with polycomb repressor complex 2 (PRC2) (De Raedt et al., 2014; Zhang et al., 2020) suggesting that epigenetic approaches to therapy may prove successful. Despite convincing evidence that epigenetic therapies including BET bromodomain inhibitors (Patel et al., 2014) and HDAC inhibitors (Lopez et al., 2011; Wojcik et al., 2019) may prove useful in patients with MPNST, clinical studies have not demonstrated successful responses to date largely due to the narrow therapeutic window of such drugs in solid tumors (Manta et al., 2022; Trojer et al., 2022) as well as acquired resistance mechanisms to such therapies (Filippakopoulos et al., 2014; Kurimchak et al., 2016; Cooper et al., 2019; Mrakovcic et al., 2019).

Our laboratory recently discovered a potent and selective inhibitor of the LHC repressor complex, corin, which has proven effective at inhibiting tumor cell growth in melanoma, cutaneous squamous cell carcinoma, diffuse intrinsic pontine glioma (DIPG) and breast cancers (Kalin et al., 2018; Anastas et al., 2019; Garcia-Martinez et al., 2022). The bifunctional nature of this inhibitor allows for targeting of both the histone deacetylase (HDAC) and lysine-specific demethylase 1 (LSD1) functions of the LHC complex and has demonstrated increased potency and an improved therapeutic window versus the corresponding Class I HDAC inhibitor (entinostat) or LSD1 inhibitor (GSK-LSD1) alone (Kalin et al., 2018; Wu et al., 2024). Additionally, corin has proven effective in inhibiting tumor cell plasticity and reversing therapy resistance in human melanoma cells (Wu et al., 2024). MPNST and melanoma tumor cells are both derived from the same embryonic neural crest cell lineage (Kiuru and Busam, 2017) and recent studies have determined that PRC2 loss in MPNST confers a dedifferentiated early neural crest phenotype (Kochat et al., 2021) similar to that seen during melanoma phenotype switching (Karras et al., 2022). As we have determined that the LHC inhibitor, corin, is able to block tumor cell growth, reverse phenotype switching and prevent establishment of neural crest-associated lineages in melanoma (Wu et al., 2024), we hypothesized that inhibition of the LHC complex in MPNST would similarly block tumor cell growth and migration, with a preference for PRC2-negative tumors. Here, we show that corin treatment of MPNST cells leads to inhibition of tumor cell growth and increased apoptosis with a significantly lower IC50 versus the HDAC inhibitor (HDACi) entinostat or the LSD1 inhibitor (LSD1i) GSK-LSD1. Interestingly, transcriptomic analysis demonstrates a specific role for the LHC complex in regulating expression of genes associated with MPNST neuronal differentiation and collagen-containing extracellular matrix without associated increases in chromatin accessibility, while corin also significantly reduces MPNST cell invasion. These data identify a specific influence of the LHC complex in regulating MPNST growth, cell death, differentiation and invasion and suggest that LHC inhibitors may prove useful in the treatment of this refractory malignancy.

## Results

### The LHC inhibitor, corin, blocks tumor cell growth and promotes apoptosis in MPNST

As previous studies have shown that inhibition of HDAC1, a component of the LHC complex, inhibits growth of MPNST cell lines harboring PRC2 complex mutations (Lopez et al., 2011), we sought to determine the effect of our bi-functional HDAC1-LSD1 inhibitor, corin, on MPNST proliferation using a panel of ten established MPNST cell lines. Eight of the cell lines evaluated possessed inactivating mutations of NF1 and the PRC2 complex, while the remaining two cell lines originating from sporadic MPNSTs possessed wildtype NF1 and PRC2 (Table 1). All MPNST cell lines were treated with either DMSO, HDACi (entinostat), LSD1 inhibitor (GSK-LSD1) or corin and evaluated in a dose-response curve (Figure 1A). Corin and entinostat treatment led to decreased proliferation across all cell lines tested without any notable response to GSK-LSD1 in any of the cell lines evaluated. Of note, the IC50 values for corin were consistently lower (1.4x-3.9x) than those observed for the HDACi, entinostat, across all cell lines (Table 2) with an IC50 range of 0.18-1.7μM for corin versus 0.4-5.0μM for entinostat. Additionally, all MPNST cells wildtype for PRC2 complex were less responsive to corin and HDACi (Figure 1A, Table 2), consistent with previously published data (Lopez et al., 2011). Annexin V staining was performed to assess the impact of corin treatment on cellular apoptosis in two NF1/PRC2 wildtype MPNST cell lines (MPNST 724/STS26T) and two NF1-/PRC2-cell lines (S462/T265) using flow cytometry. Consistent with the dose-response data (Figure 1A), corin induced apoptosis in all four cell lines tested (Figure 1B, Supplementary Figure 1) resulting in increased cell death in all cell lines treated with 1μM corin (Figure 1C). Notably, MPNST cells treated with DMSO, corin, or HDACi demonstrated significant cell death when visualized by phase contrast microscopy, with an elongated spindle-cell phenotype observed in T265 cells treated with corin (Figure 1D).

**Figure 1.**
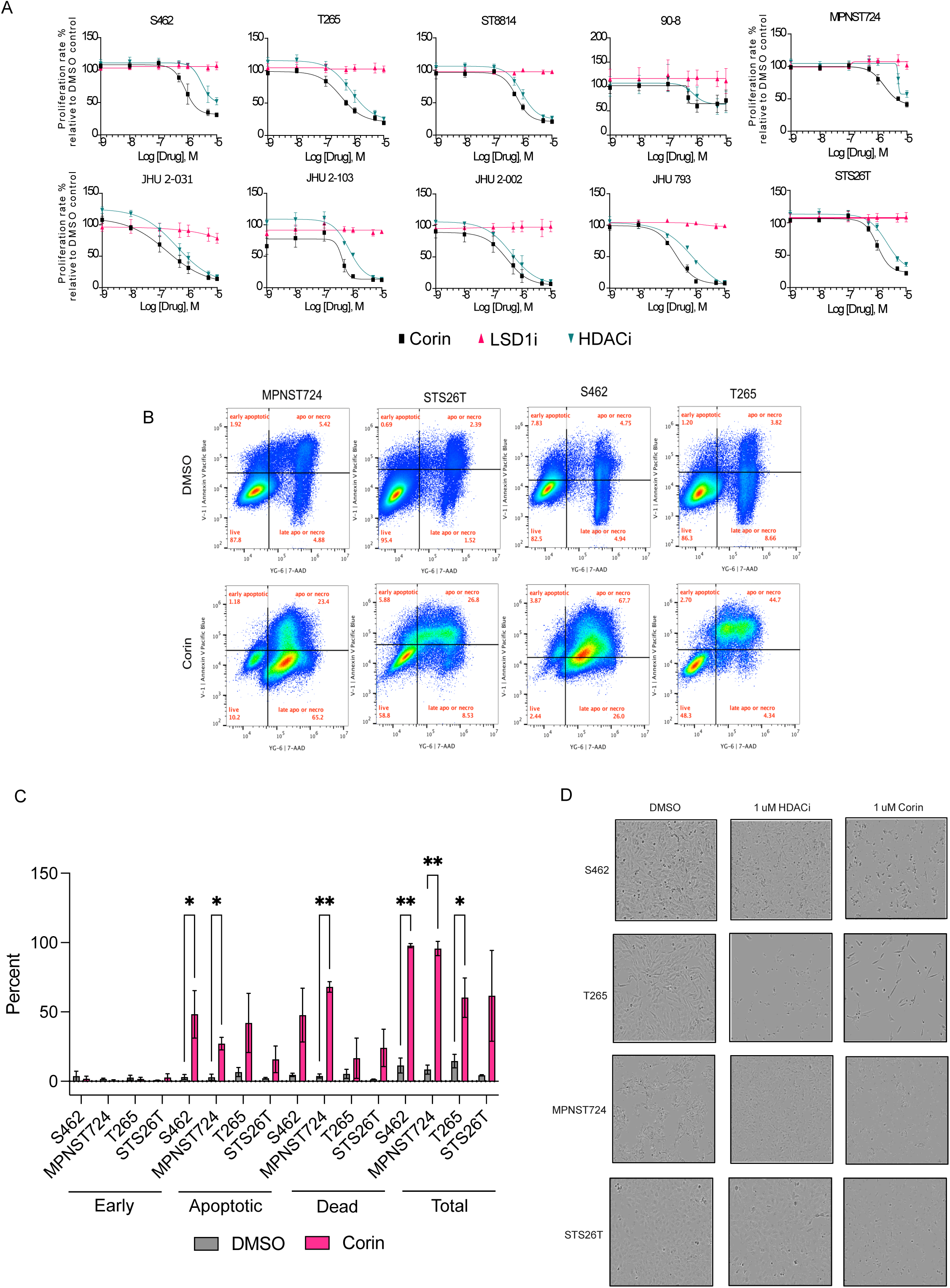
The LHC inhibitor, corin, blocks tumor cell growth and promotes apoptosis in MPNST. **A.** Dose-response curves for MPNST cells treated with DMSO, corin, HDACi (entinostat), or LSD inhibitor (GSK-LSD1). Treatment was for 72 hours except for the four JHU cell lines which were treated for 120 hours. Y axis represents proliferation rate normalized to DMSO. **B.** Flow cytometry analysis of MPNST cells treated with DMSO or 1 μM Corin for 72 hours and stained with Annexin V. **C.** Quantification of apoptosis data in **B** for early apoptotic, apoptotic/necrotic and late apoptotic/necrotic cells treated with DMSO vs. corin. **D**. Phase-contrast images of cells treated with DMSO or 1 μM Corin or HDACi for 72 hours taken at 10X magnification. Paired t test was performed for apoptosis analysis of DMSO vs. corin treatment. *P < 0.05, **P < 0.01, ***P < 0.001

**Table 1.** Characteristics of MPNST cell lines used in this study where WT indicates wildtype gene (NF1, PRC2 complex) is present and – indicates an inactivating mutation is present in the associated gene of interest.

| Cell Line | NF1 status | PRC2 status |
| --- | --- | --- |
| MPNST724 | WT | WT |
| STS26T | WT | WT |
| S462 | - | - |
| T265 | - | - |
| ST8814 | - | - |
| 90-8 | - | - |
| JH-2-031 | - | - |
| JH-2-103 | - | - |
| JH-2-079-c | - | - |
| JH-2-002 | - | - |

**Table 2.** IC50 for all ten MPNST cell lines treated with corin versus the HDACi, entinostat.

| Cell Line | Corin IC50 (uM) | HDACi IC50 (uM) |
| --- | --- | --- |
| MPNST 724 | 1.7 | 5.0 |
| STS26T | 1.0 | 2.1 |
| S462 | 0.84 | 3.0 |
| T265 | 0.34 | 0.73 |
| ST8814 | 0.60 | 0.92 |
| 90-8 | 0.49 | 0.75 |
| JH-2-031 | 0.18 | 0.40 |
| JH-2-103 | 0.42 | 0.74 |
| JH-2-079-c | 0.2 | 0.78 |
| JH-2-002 | 0.30 | 0.42 |

### Corin treatment of MPNST cells alters expression of axonogenesis and collagen-containing extracellular matrix genes independent of NF1 and PRC2 mutation status

Given the significant growth inhibitory and apoptosis-inducing effects of corin in MPNST cells, we next sought to determine the global transcriptional effects of LHC inhibition in PRC2+/NF1+ and PRC2-/NF1- MPNST cells. S462, T265, MPNST724, and STS26T cells were treated with 1uM corin for 24 hours and evaluated by RNA-seq. Remarkably, hierarchical clustering of differentially expressed genes did not group cell transcriptional profiles based on NF1/PRC2 status (Figure 2A). Principal Component Analysis (PCA) of differentially regulated genes further confirmed these findings, as MPNST cells treated with corin did not cluster according to NF1/PRC2 mutational status (Figure 2B). Given that the LHC complex primarily functions by promoting repressive histone marks (Andrés et al., 1999), it is not surprising that the vast majority of genes altered in expression following corin treatment were upregulated (Figure 2A). We therefore chose to focus our subsequent data analyses on genes that were increased in expression following corin treatment. Examination of overlapping upregulated genes from all four MPNST cell lines identified 116 common genes with the greatest number of upregulated genes seen in T265 cells and the smallest number of upregulated genes seen in MPNST724 cells (Figure 2C). When datasets from all four cell lines were combined and differential expression analysis was conducted we identified 1,435 genes that were significantly upregulated by corin in the MPNST cell lines evaluated with an additional 103 genes that were significantly downregulated by corin (Figure 2D). Further analysis of RNA-seq data from individual cell lines confirmed these results (Supplementary Figure 2). The top gene ontology (GO) pathways for genes upregulated by corin revealed significant enrichment in biological processes including axonogenesis, as well as cellular components including neuronal cell body, collagen-containing extracellular matrix, distal axon, and synaptic vesicle membrane (Figure 2E). These pathways suggest that the LHC complex influences MPNST differentiation and extracellular matrix dynamics. Consistent with this, GO analysis of differentially expressed genes in each cell line highlighted similar enriched pathways, including axonogenesis and collagen-containing extracellular matrix (Supplementary Figure 3). The top five upregulated genes within these pathways—MAPT, NGFR, SPP1, GDF15, and PDGFB were subsequently analyzed and confirmed to be upregulated in the four MPNST cell lines, (Figure 2F, 4A). MAPT is a microtubule-associated protein with tumor-suppressive properties, with knockdown in clear cell renal cell carcinoma leading to increased tumor cell growth and invasion (Han et al., 2020). NGFR is a growth factor whose expression correlates with the activation of pro-apoptotic pathways (Chen et al., 2021). SPP1 (secreted phosphoprotein-1) encodes the gene for osteopontin, an extracellular glycol phosphoprotein associated with diverse cellular functions due to a variety of distinct functional domains, proteolytically-cleaved and alternatively spliced variants, and numerous post-translational modifications (Yim et al., 2022). Although originally identified as a highly phosphorylated protein present in bone matrix, subsequent studies have determined a wide-range of expression of SPP1 in mesenchymal cells and epithelial cells, with the highest levels of expression seen in brain tissue with associated neuroprotective functions including Schwann cell-derived peripheral axon regeneration (Wright et al., 2014; Yim et al., 2022). GDF15, a member of the TGF-B superfamily, has been shown to reduce tumor cell proliferation and invasiveness and suppress epithelial-mesenchymal transition in cancer cells (Tsui et al., 2015). PDGFB, a member of the platelet-derived growth factor family, is typically seen as a homodimer (PDGF-BB) with known functions in angiogenesis (Hosaka et al., 2013). Although primarily thought of as a tumor promoting growth factor, studies suggest that functions of PDGFB may be tumor inhibitory and suppress metastasis due to specific effects on vascular remodeling and the extracellular matrix and depend on expression levels in vivo (Hosaka et al., 2013; Zhang et al., 2020; Prakash et al., 2022).

**Figure 2.**
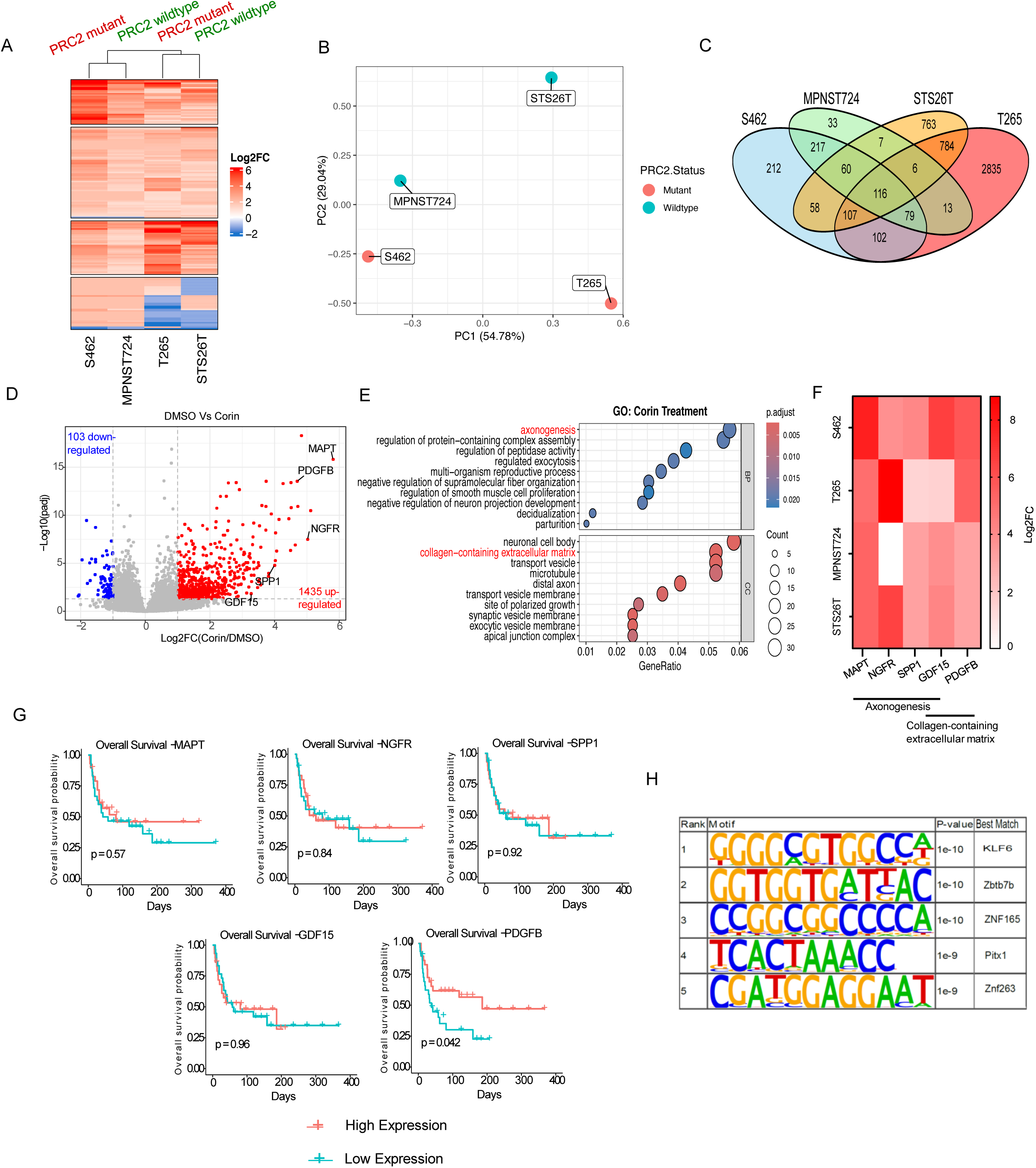
Corin treatment of MPNST cells alters expression of axonogenesis and invasion genes independent of NF1 and PRC2 mutation status. **A.** Heatmap and hierarchical clustering of genes differentially regulated by Corin in four MPNST cell lines. **B.** PCA plot for MPNST cells depicted in **A** following corin treatment. **C.** Venn diagram illustrating the overlap of differentially regulated genes among the four MPNST cell lines treated with corin. **D.** Volcano plot representing differentially expressed genes (DEGs) in all four MPNST cell lines treated with corin. Red circles indicate genes upregulated by corin, while blue circles indicate gene downregulated by corin. Genes of interests are labeled. **E.** Gene ontology analysis of the 1435 genes upregulated by corin in all four cell lines. BP: Biological process, and CC: Cellular component. **F.** Heatmap of the five genes of interests labeled based on the GO analysis. **G.** Kaplan Meier survival curves of the five genes (n=59; “high” and “low” expression defined by median expression across patients). **H.** Homer Motif analysis of the 1435 genes upregulated by Corin in all four cell lines.

In order to determine the potential functional consequences of genes upregulated in MPNST cells following corin treatment, Kaplan-Meier survival curves were generated for the upregulated genes of interest using a MPNST patient cohort (Høland et al., 2023). Among these genes, only PDGFB showed a significant positive association with survival outcome, where higher expression correlated with improved survival (Figure 2G).

We next sought to identify transcription factors associated with increased gene expression following corin treatment of MPNSTs, and performed motif analysis of the 1435 differentially upregulated genes. The top hit in our analysis was KLF6, a known tumor-suppressor transcription factor regulating cellular senescence (Figure 2H) (Narla et al., 2001; Sabatino et al., 2019). Further analysis of transcription factor motifs associated with increased gene expression following corin treatment in each cell line identified enriched motifs associated with neuronal transcription factors including TFAP2A (Hovland et al., 2022), RBPJ (Hu et al., 2011) and tumor suppressors including KLF4 (Cercek et al., 2015) and Osr1 (Yu et al., 2023) (Supplementary Figure 4).

### Inhibition of the LHC complex promotes histone acetylation without a corresponding increase in chromatin accessibility

The catalytic activity of the LHC complex is mediated by both LSD1 and HDAC1. We have previously shown that the LHC complex modulates Histone 3 Lysine 9 acetylation (H3K9ac) and Histone 3 Lysine 27 acetylation (H3K27ac) in melanoma cell lines (Wu et al., 2024) and therefore sought to determine whether corin treatment of MPNST cells would alter H3K9/H3K27 acetylation and chromatin accessibility. Four MPNST cell lines were treated with 1 µM Corin for 24 hours and H3K9ac and H3K27ac levels were evaluated by western blot. Interestingly, an increase in H3K27ac was only seen in S462 cells, while other cell lines showed an increase in H3K9ac (Figure 3A, 3B) with the greatest increases noted in S462 cells. In order to evaluate chromatin accessibility changes associated with corin treatment of MPNST, S462 cells treated with DMSO, 1 µM corin, or 1 µM entinostat for 24 hours were evaluated by ATAC-seq. Surprisingly, both corin and HDACi treatments led to a decrease in chromatin accessibility in S462 cells (Figure 3C) despite the well-documented chromatin repressor activities of LHC and HDACs. Annotation of the accessibility peaks across the genome revealed no notable differences between cells treated with DMSO, corin or the HDACi, entinostat (Figure 3D). Additionally, visualizing individual gene tracks for the five genes identified in our RNA-seq experiments (Figure 2F) revealed no significant differences in chromatin accessibility (Figure 3E).

**Figure 3.**
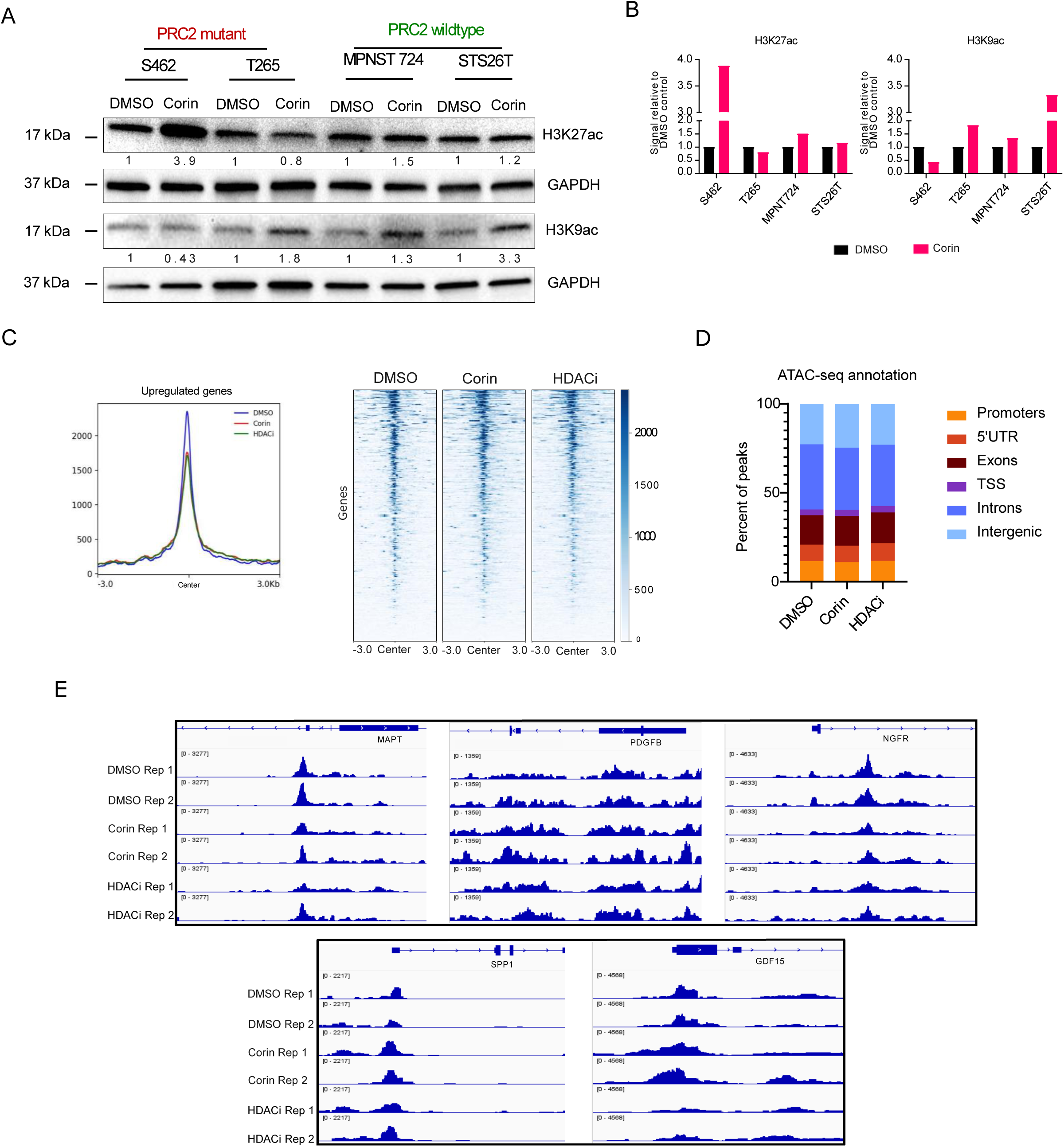
Inhibition of the LHC complex promotes histone acetylation without a corresponding increase in chromatin accessibility. **A.** Western blot of histone marks in PRC2+ (wildtype) and PRC2- (mutant) MPNST cells treated with DMSO vs corin. **B.** Bar graph quantifying the western blot in **A. C.** Plot profile and heatmap of chromatin accessibility in S462 MPNST cells treat with DMSO, corin or HDACi (entinostat). X axis represents the center of peak spanning −3k to 3k bp. Y axis represents the accessibility signal. **D.** Genome annotation illustrating alteration of accessibility peaks at different genome annotations in S462 cells following treatment with DMSO, corin, or HDACi (entinostat). **E.** Individual chromatin accessibility gene tracks for MAPT, PDGFB, NGFR, SPP1 and GDF15 in S462 cells following treatment with DMSO, corin or HDACi (entinostat).

### Inhibition of the LHC complex leads to decreased tumor cell invasion in MPNST

RNA-seq data identified GO pathways associated with axonal differentiation and extracellular matrix pathways as being significantly enriched in corin-treated MPNST cells (Figure 2). As these pathways are known to influence tumor cell metastatic phenotypes, we therefore investigated whether corin treatment of MPNST cells could impact tumor cell invasion. Five genes associated with neuronal differentiation or extracellular matrix composition which were identified as being significantly upregulated by corin in our RNA-seq studies were validated using qPCR (Figure 4A) with all four cell lines demonstrating increased expression of the genes of interest following corin treatment. Boyden Chamber invasion assays were subsequently performed on all four MPNST cell lines to assess changes in MPNST invasion following 24 hour treatment with 1 µM corin (Figure 4B, 4C). All MPNST cell lines demonstrated reduced invasion following corin treatment with the most highly invasive cells lines in the DMSO controls (S462, STS26T) demonstrating the most significant decreases in invasiveness in response to corin. Overall, these findings suggest a critical role for the LHC repressor complex in mediating important tumorigenic functions in MPNST cells with LHC inhibition leading to decreased cell proliferation, increased apoptosis, increased cellular differentiation and decreased tumor cell invasion regardless of PRC2 status (Figure 4D).

**Figure 4.**
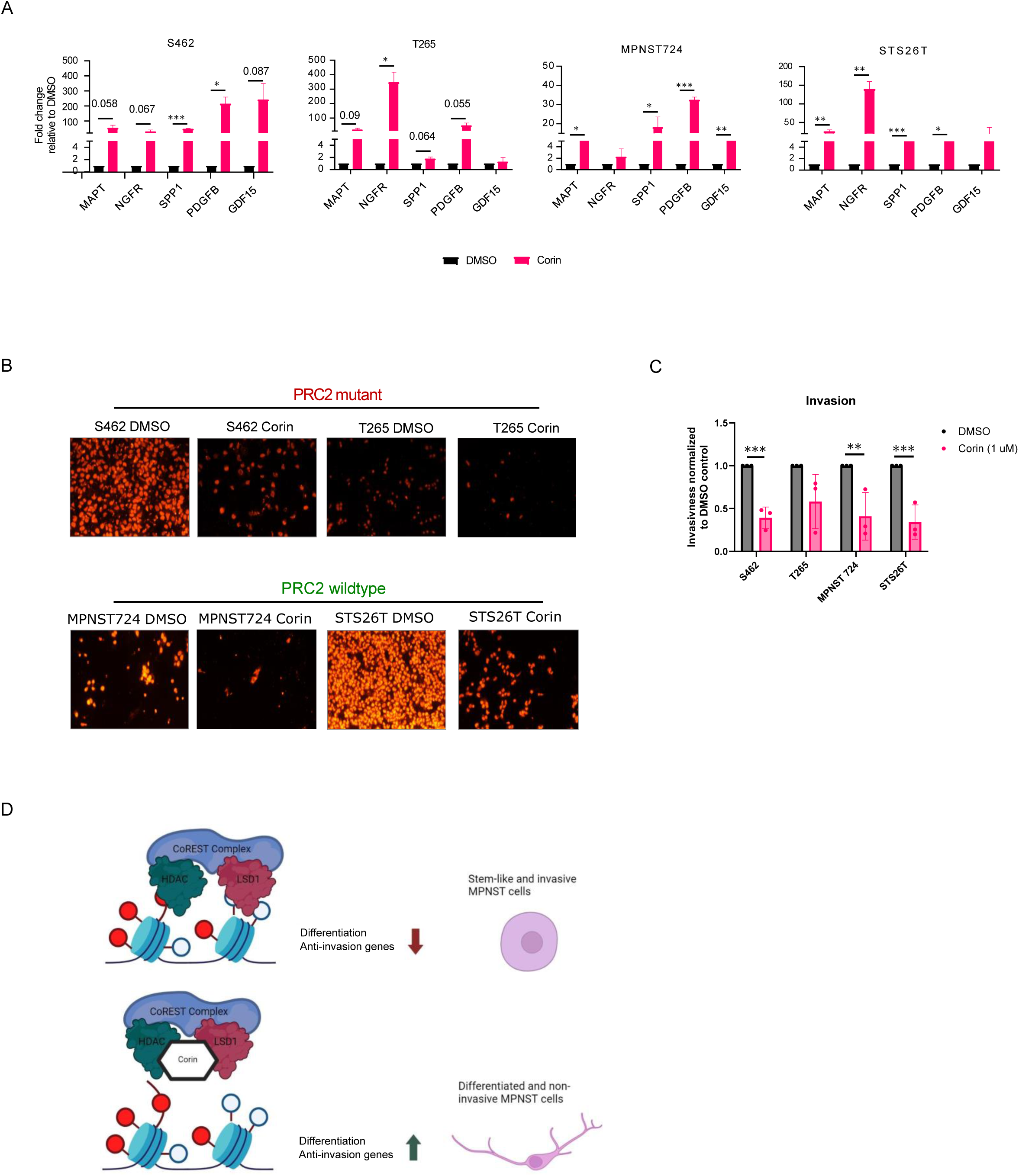
Inhibition of the LHC complex leads to decreased tumor cell invasion in MPNST. **A.** qPCR analysis of expression of genes of interest in S462, T265, MPNST724 and STS26T MPNST cells following treatment with DMSO or 1μM corin for 24 hours. **B.** Boyden chamber invasion assay of S462, T265, MPNST724 and STS26T MPNST cells treated with DMSO or 1 μM corin. **C.** Quantification of Boyden chamber invasion assay in S462, T265, MPNST724 and STS26T MPNST cells treated with DMSO or 1 μM corin. **D.** Working model depicting the repressive function of the CoREST complex on differentiation and invasion genes in MPNST cells, leading to a stem-like cell that is invasive. Treatment with corin leads to the depression of differentiation and invasion genes, leading to a more differentiated cell that is less invasive.

## Discussion

In this study, we explore the role of the CoREST epigenetic repressor complex in regulating tumorigenesis in MPNST. Using the bifunctional small molecular inhibitor of LHC, corin, we demonstrate that the catalytic subunits of the CoREST complex are critical mediators of tumor cell proliferation, apoptosis and invasion of MPNST cells. Inhibition of the LHC complex by corin leads to a significant reduction in cell proliferation, with significantly greater effects than the HDACi, entinostat in all MPNST cell lines evaluated; however, NF1+/ PRC2+ tumor cells demonstrate decreased sensitivity to corin versus NF1-/PCR2-tumor cells (Figure 1A and Table 2). Additionally, we find that corin induces apoptosis in MPNST cell lines regardless of their PRC2 mutational status (Figure 1B); this is in contrast to the relative resistance of PRC2+ MPNST cells to apoptosis in the setting of treatment with entinostat and other HDACi therapies (Hirokawa et al., 2005; Lopez et al., 2011) Interestingly, previous studies have shown that the HDAC8-specific inhibitors PCI3 and PIC4 inhibit MPNST cell growth irrespective of PRC2 mutation status (Lopez et al., 2015); however, the IC50 for these reagents is 5-10-fold that of corin for a 96-hour treatment regimen suggesting lower sensitivity to these reagents.

Transcriptomic analysis of corin-treated MPNST cells revealed significant upregulation of gene expression in all cell lines, with relatively few genes downregulated in response to corin (Figure 2A, 2D) which is consistent with the LHC complex functioning as a transcriptional repressor complex and previous data in other cells (Kalin et al., 2018; Anastas et al., 2019; Garcia-Martinez et al., 2022; Wu et al., 2024; Almier et al., 2024). Remarkably, we found top gene signatures upregulated in all MPNST cells treated with corin to include biological processes and cellular components associated with neuronal differentiation and axonogenesis which is consistent with CoREST complex functions as inhibitors of neuronal differentiation (Sáez et al., 2015). In addition, collagen-containing extracellular matrix pathways were among the top enriched GO signatures in all MPSNT cells treated with corin (Figure 2E, Supplementary Figure 2). One of the most upregulated genes in the collagen-containing extracellular matrix pathway was PDGFB (Figure 2D, 2F), and Kaplan-Meier survival data indicated a significant survival advantage associated with higher PDGFB expression in MPNST. This was particularly surprising, given the known functions of PDGFB as a growth factor for mesenchymal cells; however, within the context of extracellular matrix remodeling, PDGFB may be an important mediator of the antitumor effects of corin in MPNST (Zhang et al., 2020) which could account for corin-associated inhibition of tumor cell invasion (Figure 4B, 4C). Consistent with the Gene Ontology (GO) data, transcription factor motifs associated with promoter regions of upregulated genes following corin treatment included binding sites associated with neuronal differentiation and tumor suppression.

Unexpectedly, ATAC-seq analysis of S462 cells demonstrated that genome-wide chromatin accessibility was reduced in these PRC2-mutant MPNST cells following treatment with corin or the HDACi, MS-275 (Figure 3C), despite increases in H3K9/H3K27 acetylation (Figure 3A, 3B) suggesting a more open chromatin structure (Klemm et al., 2019). Notably, basal chromatin accessibility was found to be high both genome-wide and at specific gene sites (Figure 3C and 3E) in these PRC2-negative cells, which is not surprising given the repressor functions of the PRC2 complex (Liu and Liu, 2022) which are lost in these cells. These data suggest that the primary function of corin in these tumor cells may involve altered transcription factor recruitment to chromatin rather than chromatin accessibility which is consistent with a recent report suggesting that increases in chromatin accessibility within open chromatin regions does not necessarily correlate with gene expression changes (Kiani et al., 2022).

Finally, we confirmed the upregulated expression of several genes associated with axonogenesis/neuronal differentiation and extracellular matrix remodeling in MPNST cells treated with corin and found that corin potently inhibits tumor cell invasion in MPNST cells irrespective of PRC2 status. Interestingly, recent studies using isogenic murine MPNST cell lines show that PRC2 loss in MPNST is a critical driver of tumor progression which leads to increased expression of matrix metalloproteinases (MMPs) and other matrix remodeling enzymes as well as increased collagen-dependent invasion and metastasis (Brockman et al., 2022). Thus, the anti-invasive properties associated with corin treatment of MPNST cells may be particularly useful in PRC2 inactivated tumors.

MPNST is a highly aggressive tumor and the leading cause of death in patients with Neurofibromatosis Type 1. Despite significant advances in targeted and immunotherapy treatment of solid tumors over the past 3 decades, the prognosis for patients with MPNST has not changed substantially during this time and there are currently no FDA-approved systemic therapies for this aggressive malignancy (reviewed in (Somatilaka et al., 2022)). Significant data suggests that epigenetic approaches to treat MPNST are warranted; however, no epigenetic therapies have proven successful in altering patient outcomes to date. Here, we show that the CoREST complex is a specific vulnerability in MPNST, particularly in PRC2-null tumors, and that the potent and specific bifunctional CoREST inhibitor, corin, demonstrates significant effects in inhibiting tumor cell growth, inducing apoptosis, promoting tumor cell differentiation and inhibiting tumor cell invasion in MPNST. Given the poor prognosis of this aggressive tumor with limited treatment options for advanced stages of disease, we suggest that further preclinical studies are warranted to explore the therapeutic potential of corin in patients with MPNST.

## Materials and Methods

### Cell Culture

The S462 and MPNST724 cell lines were graciously provided by Dr. Raphael Pollock. Additionally, the ST88-14, T265, 90-8, and STS26T cell lines were generously shared by Dr. Jeffrey Field. The JH-2-002, JH-2-031, JH-2-079c, and JH-2-103 patient-derived MPNST cell lines (Wang et al., 2020) originated from the Johns Hopkins NF1 biospecimen repository (Banerjee et al., 2024; Pollard et al., 2020) and were shared under Material Transfer Agreement from Johns Hopkins University. The 6 non-JHU affiliated cell lines were cultured in high glucose DMEM with pyruvate (Thermo Fisher Scientific) supplemented with 10% Fetal Bovine Serum (FBS) and 1% penicillin/streptomycin. The JH patient-derived cell lines were cultured in high glucose DMEM (Thermo Fisher Scientific) supplemented with 10% FBS and 1% penicillin/streptomycin. All cell lines were maintained at 37°C and 5% CO_2_.

### Compounds

The HDAC inhibitor Entinostat/MS275 (Cat. No.: HY-12163) and LSD1 inhibitor GSK-LSD1 (Cat. No.: HY-100546) were purchased from MedChemExpress. Corin was prepared as previously described (Kalin et al., 2018). All stock solutions were dissolved in DMSO.

### Quanti-iT PicoGreen dsDNA Cell Proliferation Assay

Cells were seeded in a 96-well and treated with the respective compound 24 hours later. PicoGreen assay was done according to the manufacturer’s suggested protocol (Thermo Fisher Scientific).

### Apoptosis assay

Apoptosis was measured using the Pacific Blue™ Annexin V/SYTOX™ AADvanced™ Apoptosis Kit (ThermoFisher, A35136). The experimental protocol was performed according to the manufacturer’s recommendations. Spectral flow analysis was performed using Cytek Aurora (Cytek Biosciences) according to the manufacturer’s recommendations. Flow cytometry data was analyzed using SpectroFlo provided by Cytek Biosciences and FlowJo software.

### RNA extraction

S462, T265, MPNST724, and STS26T cells were treated for 24 hours with 1 μM Corin or DMSO control treatment. RNA was extracted using the RNeasy Plus Mini Kit (Qiagen).

### RNA-seq

Library was prepared and sequenced by Azenta Life Sciences. Reads were trimmed and low-quality reads were removed using Trimmomatic v.0.36. Reads were mapped to the GRCh38 reference genome using STAR aligner v.2.5.2b. Unique gene hit counts were calculated by using featureCounts from the Subread package v.1.5.2. Counts were used for differential analysis using DESeq2. The Wald test was used to generate p-values and log2 fold changes. Genes with an adjusted p-value < 0.05 and absolute log2 fold change > 1 were called as differentially expressed genes for each comparison. Heatmap and hierarchical clustering for the RNA-seq was generated using ComplexHeatmap v.2.20.0 (Gu et al., 2016). PCA analysis was done using ggfortify v.4.4.1. Volcano plots were generated using ggplot2 and labeling was done using ggrepel. GO Plots were generated using the enrichGO and dotplot commands in clusterProfiler v4.12.0 (Wu et al., 2021). Motif analysis was done using Samtools v.1.12 and homer v.4.1 using the findmotif.pl command searching for motifs within −400 to 100 bp of TSS.

### Kaplan Meier survival curves

Kaplan Meier plots were generated using patient outcome and expression data (Høland et al., 2023). Patients were excluded if mRNA expression analysis was unavailable, the patient died from other causes, or lacked follow-up. Plots and statistical analysis were performed in R using survfit from the survival package (v2.11-4).

### Whole Cell Extracts and SDS-PAGE

Proteins were extracted from cells using M-PER lysis buffer according to the manufacturer’s recommendations (Thermo Fisher Scientific). Protein concentrations were determined using the Pierce™ BCA Protein Assay (Thermo Fisher Scientific). 10 ug of whole cell extracts were separated on gradient 4-20% polyacrylamide precast gels (Bio-Rad) and transferred to PVDF membranes (Bio-Rad). Membranes were incubated with the following antibodies overnight: H3K9ac (ab32129), H3K27ac (ab4729), or GAPDH (sc-365062).

### ATAC-seq

100,000 cells were seeded and treated with 1 μM Corin or DMSO. DNA was extracted, tagmented, purified, and adapter ligated using the ATAC-seq Kit (53150) according to the manufacturer’s instructions (Active Motif). Sequencing was done by Azenta Life Sciences and reads were trimmed using the fastp command in fastp v.0.23.4. Reads were aligned with GRCh38 reference genome using bowtie2 v.2.4.2. Sam files were filtered to retain the mapped reads only and converted to BAM files using Samtools v.1.12. Bed and Bedgraph files were created using Bedtools v.2.31.0. Peaks were called using Macs2 v.2.2.7.1 with the nomodel, and nolambda options. Bam files were normalized using RPKM to create BigWig files using the bamCoverage command in deeptools v.3.5.1. Deeptools was also used to create the matrix using the command computeMatrix, prolfile was created using the command plotProfile, and heatmaps were created using the command plotHeatmap. BigWig files were used to visualize individual gene tracks using IGV v.2.17.4. Genome annotation was done using Samtools v.1.12 and homer v.4.1 with the command annotatePeaks.pl.

### cDNA preparation and qPCR

1 μg of RNA was reverse transcribed using the SuperScript III First-Strand Synthesis System kit (Invitrogen, Thermo Fisher Scientific). 40 cycles of qPCR were performed using the Step One Plus Real-time PCR System (Applied Biosystems). Data represents 3 biological replicates with 3 technical replicates per experiment. Primers used are listed in Supplementary Table 1.

### Boyden chamber invasion assay

Transwell inserts (8 μm pore membrane) were placed in each well of a 24-well plate. 50 μg of Matrigel (Corning) diluted in 30 μL serum-free DMEM was added to each insert and incubated at 37 °C for 30 minutes. 150,000 cells were seeded in 300 μL serum-free DMEM media on the Matrigel in the top chamber and treated with 1μM Corin or DMSO. 600 μL of 20% FBS DMEM was used as the attractant in the bottom chamber. After 24 hours, cells and the Matrigel in the top chamber were removed, the membrane was washed with Phosphate-buffered saline (PBS), and fixed in 70% ethanol. The bottom of the membrane (where invading cells are located) was stained using 50 μg/mL propidium iodide and washed in PBS before cutting out the membrane.

Stained membrane was visualized using the Nikon Eclipse E400 microscope and SPOT Advanced software. Five images at 20X were captured for each membrane. Image analysis was done using Fiji ImageJ using the method previously described (Schroeder et al., 2021). A total of three biological replicates were done with three technical replicates done for each experiment.

## Data and code availability

All the sequencing data generated are deposited in the GEO database (accession number GSE254326). Any additional information requested can be directed to the lead contact. No codes were generated in the manuscript.

## Acknowledgments

R.A. and M.C. are supported by a Department of Defense Translational Research Award in Melanoma (W81XWH-21-1-0980); P.A.C. is supported by NIH grant R35 GM149229. We thank Dr. Anna Belkina for her expertise and assistance with FACS protocols and data analysis. We thank Dr. Raphael Pollock for providing S462 and MPNST724 cell lines. We thank Dr. Jefferey Field for providing ST88-14, T265, 90-8, and STS26T cell lines. We thank Drs. Stavriani Makri and Christine Pratilas for providing JH-2-002, JH-2-031, JH-2-079c, and JH-2-103 cell lines.

## Author Contributions

Conceptualization: RMA; Experimental Design/Data Curation: IS, RF, SB, MC, RMA Formal Analysis: IS, RF, MC, RMA, Supervision: MC, PAC, RMA; Writing – Original Draft: IS, RMA; Writing – Review and Editing: IS, RF, SB, MC, PAC, RMA

