## Supplementary Data and Table for "The CoREST complex is a therapeutic vulnerability in malignant peripheral nerve sheath tumors"

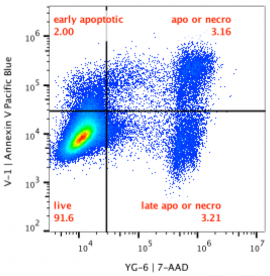


MPNST-Annexin set 5-S462 DMSO.fcs
doublet exclusion
15-Jul-2024

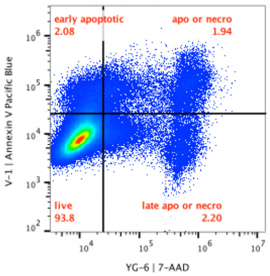


MPNST-Annexin set 5-MPNST724 DMSO.fcs
doublet exclusion
15-Jul-2024

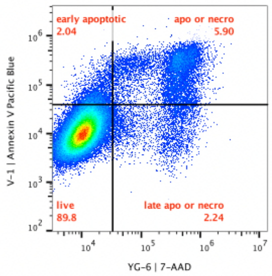


MPNST-Annexin set 5-T265 DMSO.fcs
doublet exclusion
15-Jul-2024

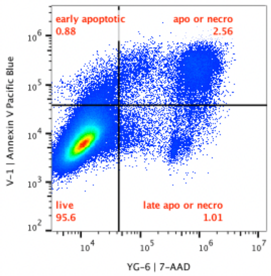


MPNST-Annexin set 5-STS26T DMSO.fcs
doublet exclusion
15-Jul-2024

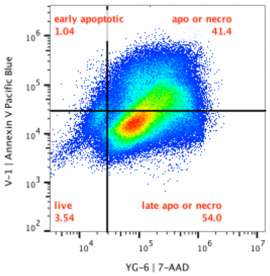


MPNST-Annexin set 5-S462 Corin.fcs
doublet exclusion
15-Jul-2024

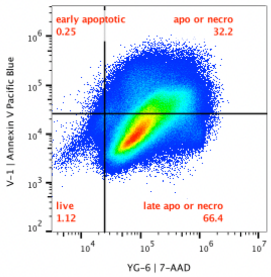


MPNST-Annexin set 5-MPNST724 Corin.fcs
doublet exclusion
15-Jul-2024

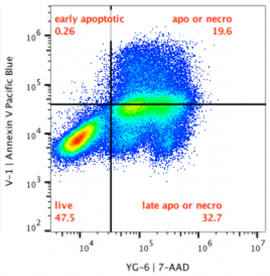


MPNST-Annexin set 5-T265 Corin.fcs
doublet exclusion
15-Jul-2024

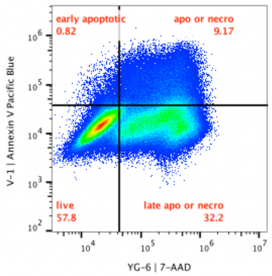


MPNST-Annexin set 5-STS26T Corin.fcs
doublet exclusion
15-Jul-2024

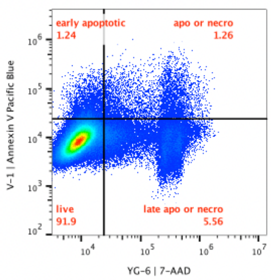


MPNST-Annexin set 6-S462 DMSO.fcs
doublet exclusion
17-Jul-2024

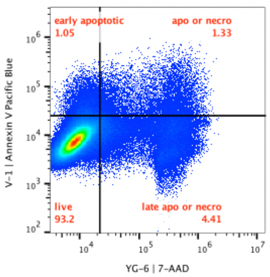


MPNST-Annexin set 6-MPNST724 DMSO.fcs
doublet exclusion
17-Jul-2024

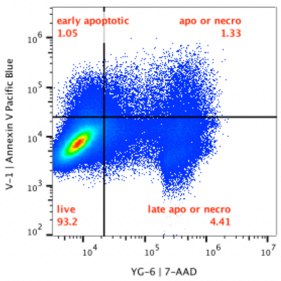


MPNST-Annexin set 6-MPNST724 DMSO.fcs
doublet exclusion
17-Jul-2024

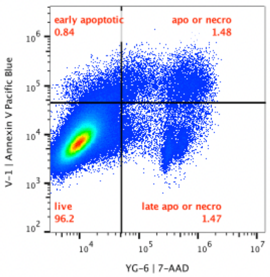


MPNST-Annexin set 6-STS26T DMSO.fcs
doublet exclusion
17-Jul-2024

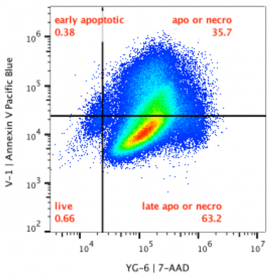


MPNST-Annexin set 6-S462 Corin.fcs
doublet exclusion
17-Jul-2024

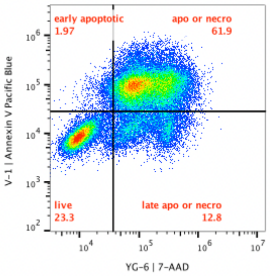


MPNST-Annexin set 6-T265 Corin.fcs
doublet exclusion
17-Jul-2024

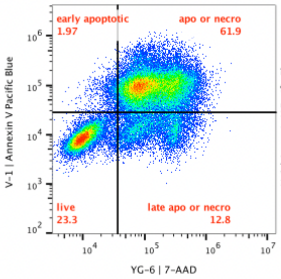


MPNST-Annexin set 6-T265 Corin.fcs
doublet exclusion
17-Jul-2024

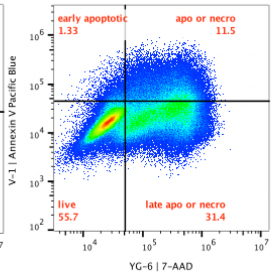


MPNST-Annexin set 6-STS26T Corin.fcs
doublet exclusion
17-Jul-2024

**Supplementary Figure 1.** Flow cytometry analysis of apoptosis using annexin V and 7AAD. Both biological replicates were used in addition to Figure 1C for the quantification and statistical analysis depicted in Figure 1D.

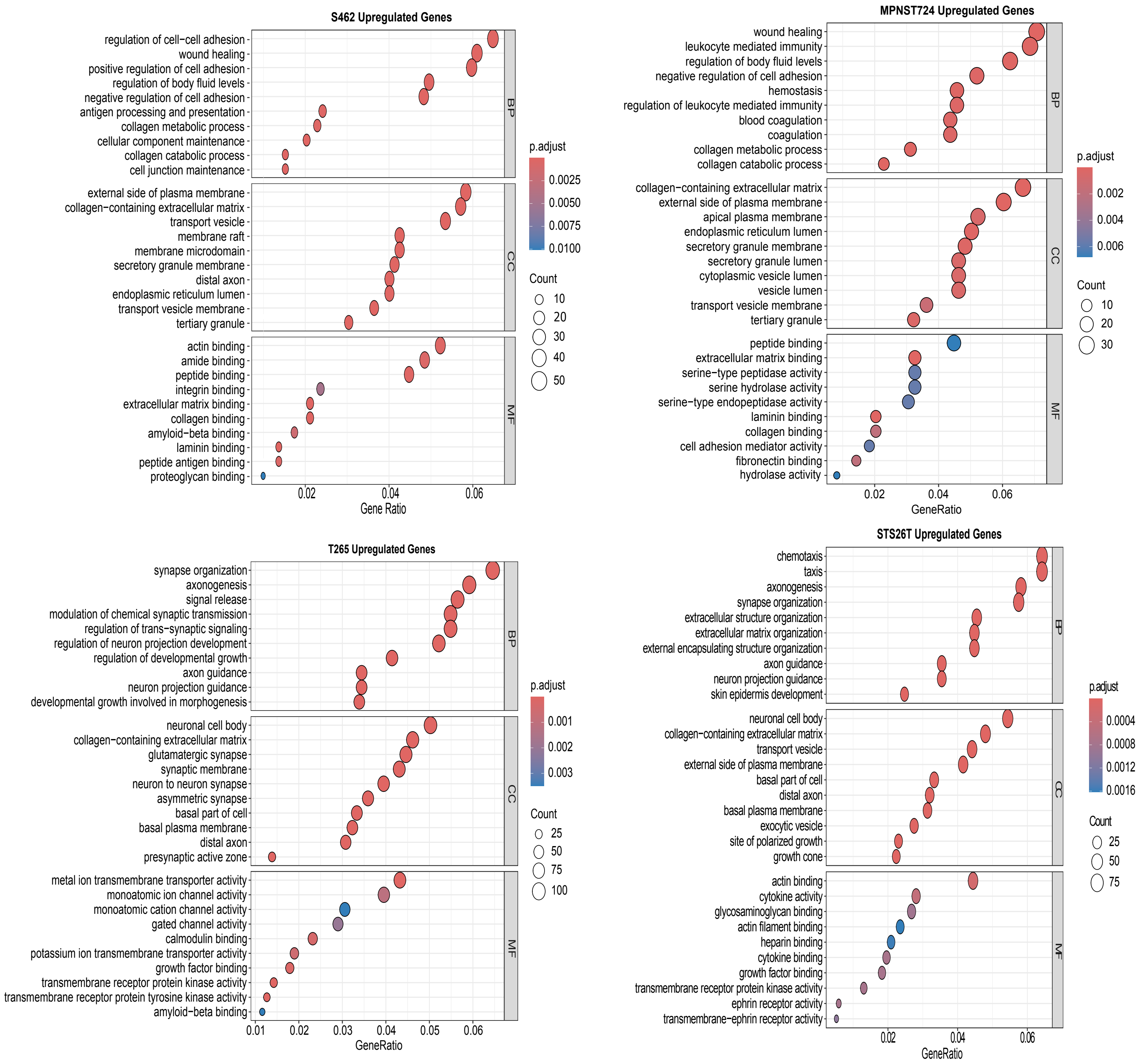
S462 Upregulated Genes
regulation of cell−cell adhesion
wound healing
positive regulation of cell adhesion
regulation of body fluid levels
negative regulation of cell adhesion
antigen processing and presentation
collagen metabolic process
cellular component maintenance
collagen catabolic process
cell junction maintenance
external side of plasma membrane
collagen−containing extracellular matrix
transport vesicle
membrane raft
membrane microdomain
secretory granule membrane
distal axon
endoplasmic reticulum lumen
transport vesicle membrane
tertiary granule
actin binding
amide binding
peptide binding
integrin binding
extracellular matrix binding
collagen binding
amyloid−beta binding
laminin binding
peptide antigen binding
proteoglycan binding
BP
CC
MF
0.02
0.04
0.06
Gene Ratio
p.adjust
0.0025
0.0050
0.0075
0.0100
Count
10
20
30
40
50
MPNST724 Upregulated Genes
wound healing
leukocyte mediated immunity
regulation of body fluid levels
negative regulation of cell adhesion
hemostasis
regulation of leukocyte mediated immunity
blood coagulation
coagulation
collagen metabolic process
collagen catabolic process
collagen−containing extracellular matrix
external side of plasma membrane
apical plasma membrane
endoplasmic reticulum lumen
secretory granule membrane
secretory granule lumen
cytoplasmic vesicle lumen
vesicle lumen
transport vesicle membrane
tertiary granule
peptide binding
extracellular matrix binding
serine−type peptidase activity
serine hydrolase activity
serine−type endopeptidase activity
laminin binding
collagen binding
cell adhesion mediator activity
fibronectin binding
hydrolase activity
BP
CC
MF
0.02
0.04
0.06
GeneRatio
p.adjust
0.002
0.004
0.006
Count
10
20
30
T265 Upregulated Genes
synapse organization
axonogenesis
signal release
modulation of chemical synaptic transmission
regulation of trans−synaptic signaling
regulation of neuron projection development
regulation of developmental growth
axon guidance
neuron projection guidance
developmental growth involved in morphogenesis
neuronal cell body
collagen−containing extracellular matrix
glutamatergic synapse
synaptic membrane
neuron to neuron synapse
asymmetric synapse
basal part of cell
basal plasma membrane
distal axon
presynaptic active zone
metal ion transmembrane transporter activity
monoatomic ion channel activity
monoatomic cation channel activity
gated channel activity
calmodulin binding
potassium ion transmembrane transporter activity
growth factor binding
transmembrane receptor protein kinase activity
transmembrane receptor protein tyrosine kinase activity
amyloid−beta binding
BP
CC
MF
0.01
0.02
0.03
0.04
0.05
0.06
GeneRatio
p.adjust
0.001
0.002
0.003
Count
25
50
75
100
STS26T Upregulated Genes
chemotaxis
taxis
axonogenesis
synapse organization
extracellular structure organization
extracellular matrix organization
external encapsulating structure organization
axon guidance
neuron projection guidance
skin epidermis development
neuronal cell body
collagen−containing extracellular matrix
transport vesicle
external side of plasma membrane
basal part of cell
distal axon
basal plasma membrane
exocytic vesicle
site of polarized growth
growth cone
actin binding
cytokine activity
glycosaminoglycan binding
actin filament binding
heparin binding
cytokine binding
growth factor binding
transmembrane receptor protein kinase activity
ephrin receptor activity
transmembrane−ephrin receptor activity
BP
CC
MF
0.02
0.04
0.06
GeneRatio
p.adjust
0.0004
0.0008
0.0012
0.0016
Count
25
50
75

**Supplementary Figure 2.** Gene Ontology (GO) analysis of differentially expressed genes in S462, T265, MPNST724 and STS26T MPNST cells treated with corin. BP: Biological process, CC: Cellular component, and MF: Molecular function.

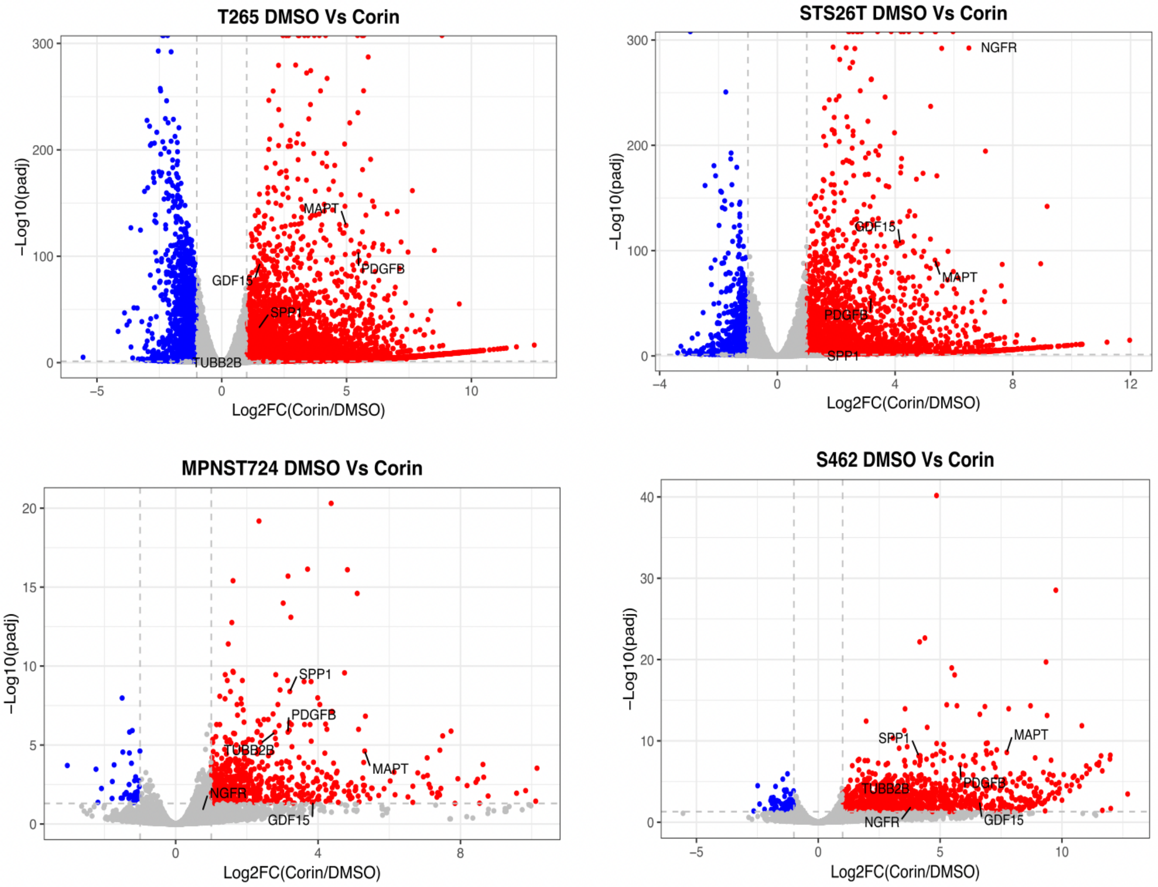

T265 DMSO Vs Corin
−Log10(padj)
Log2FC(Corin/DMSO)
MAPT
PDGFB
GDF15
SPP1
TUBB2B
STS26T DMSO Vs Corin
NGFR
GDF15
MAPT
PDGFB
SPP1
MPNST724 DMSO Vs Corin
SPP1
PDGFB
TUBB2B
MAPT
NGFR
GDF15
S462 DMSO Vs Corin
SPP1
MAPT
TUBB2B
PDGFB
NGFR
GDF15

**Supplementary Figure 3.** Volcano plot representing differentially expressed genes in S462, T265, MPNST724 and STS26T MPNST cells treated with corin. Red circles indicate genes upregulated by Corin, while blue circles indicate gene downregulated by Corin. Genes of interests are labeled.

## S462

| Rank | Motif | P-value | Best Match |
|---|---|---|---|
| 1 | 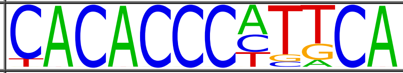 | 1e-15 | KLF4 |
| 2 | 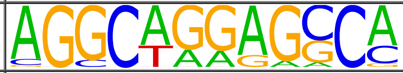 | 1e-14 | SD0002.1 |
| 3 | 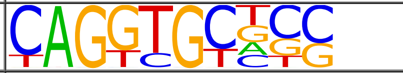 | 1e-12 | TCF4 |
| 4 | 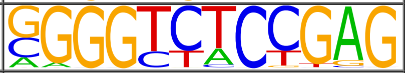 | 1e-11 | LRF(Zf) |
| 5 | 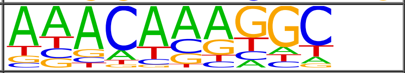 | 1e-11 | Sox3 |
| 6 | 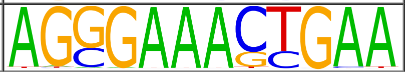 | 1e-11 | ISRE(IRF) |

## T265

| Rank | Motif | P-value | Best Match |
|---|---|---|---|
| 1 | 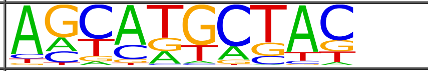 | 1e-14 | Osr1 |
| 2 | 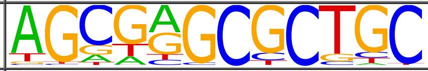 | 1e-14 | Zfp161 |
| 3 | 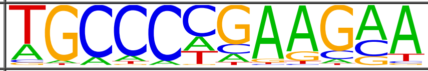 | 1e-13 | THAP1 |
| 4 | 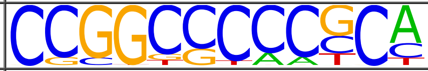 | 1e-12 | Egr1 |
| 5 | 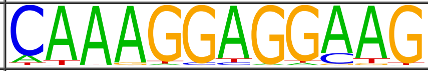 | 1e-11 | SPIC |
| 6 | 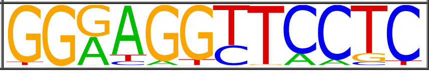 | 1e-11 | NFkB-p65-Rel |

### MPNST724

| Rank | Motif | P-value | Best Match |
|---|---|---|---|
| 1 | 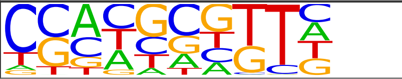 | 1e-11 | PRDM15 |
| 2 | 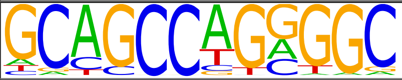 | 1e-11 | TFAP2A |
| 3 | 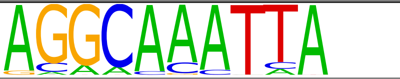 | 1e-11 | Pou2f2 |
| 4 | 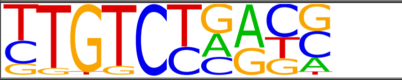 | 1e-11 | Smad3 |
| 5 | 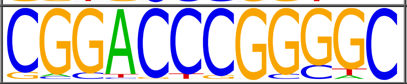 | 1e-11 | Nr2f2 |
| 6 | 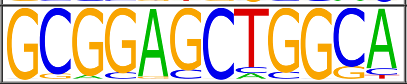 | 1e-10 | MED-1 |

### STS26T

| Rank | Motif | P-value | Best Match |
|---|---|---|---|
| 1 | 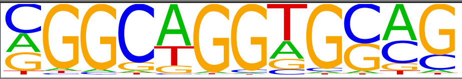 | 1e-14 | Slug |
| 2 | 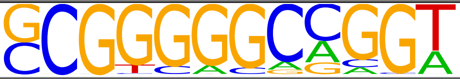 | 1e-14 | Egr1_1 |
| 3 | 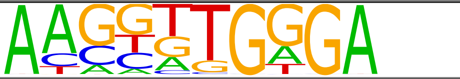 | 1e-13 | RBPJ |
| 4 | 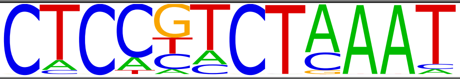 | 1e-12 | HOXB13 |
| 5 | 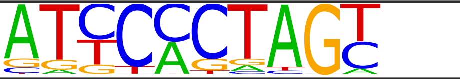 | 1e-12 | Hmbox1 |
| 6 | 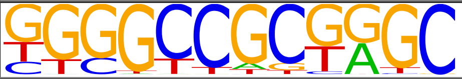 | 1e-12 | Tcfap2b |

**Supplementary Figure 4.** Homer Motif analysis of the 1435 genes upregulated by corin in S462, T265, MPNST724 and STS26T MPNST cells.

| Gene | Forwar Primer (5' -> 3') | Reverse Primer (5' -> 3') |
|---|---|---|
| MAPT | AGCCAAGACATCCACACGTT | AGAGGGTCTGAGCTACCAGG |
| NGFR | CTGGCACCGCCTTCTCTAAA | GGCAAAGCTGACTTGGCTTC |
| SPP1 | AGGCATCACCTGTGCCATAC | GGCCACAGCATCTGGGTATT |
| GDF15 | GCAAGAACTCAGGACGGTGA | TGGAGTCTTCGGAGTGCAAC |
| PDGFB | GCGCCCATTTTTCATTCCCTAGATA | GGTTTTCTCTTTGCAGCGAGGC |

**Supplementary Table 1.** Primers used for qPCR analysis of MAPT, NGFR, SPP1, GDF15 and PDGFB expression in MPNST cell lines following corin treatment.
